# Pleiotropy of Bcl-2 family proteins is an ancient trait in the metazoan evolution

**DOI:** 10.1101/816322

**Authors:** Nikolay Popgeorgiev, Lea Jabbour, Trang Thi Minh Nguyen, Nikola Ralchev, Rudy Gadet, Stéphen Manon, Hans-Jürgen Osigus, Bernd Schierwater, Ruth Rimokh, Germain Gillet

## Abstract

In the animal kingdom, proteins of the Bcl-2 family are widely recognized as regulators of mitochondrial outer membrane permeabilization (MOMP), leading to apoptotic cell death. These proteins were recently also shown to control IP_3_-dependent calcium fluxes at the level of the endoplasmic reticulum (ER). However, the origin and evolution of these pleiotropic functions remain elusive. Here, we molecularly characterized the four members of the Bcl-2 family (trBcl-2L1 to -2L4) in the most primitive metazoan, namely *Trichoplax adhaerens*. Primary structure and phylogenetic analyses demonstrated that all four trBcl-2 homologs belong to the multidomain Bcl-2 group and presented a conserved C-terminus transmembrane (TM) domain. TrBcl-2L1 and trBcl-2L2 are highly divergent proteins clustering with the anti-apoptotic Bcl-2 members, whereas trBcl-2L3 and trBcl-2L4 were homologous to the pro-apoptotic Bax (trBax) and Bak (trBak). Interestingly, at the functional level, trBak operates as a BH3 only sensitizer repressing the anti-apoptotic activities of trBcl-2L1 and trBcl-2L2, whereas trBax leads to MOMP, similarly to the well-known indirect model of Bax activation. Finally, we found that trBcl-2L1 had a dual ER and mitochondrial subcellular localization and was able to bind to IP_3_R. By generating two TM domain mutants we demonstrated that trBcl-2L1 targeted to the ER was able to control IP_3_-dependent calcium fluxes, whereas Mito-trBcl-2L1 represses trBax-dependent MOMP, suggesting that Bcl-2 pleiotropy appeared early and was conserved throughout metazoan evolution.

## INTRODUCTION

Apoptosis represents one of the hallmarks of multicellularity in the animal kingdom ^1^. This process shapes the body during the embryonic development, maintains tissue homeostasis and eliminates undesirable or potentially harmful cells from the organism ^2^. Proteins belonging to the Bcl-2 family represent one of the main regulators of apoptosis. They control a key step in the induction of apoptosis called the mitochondrial outer membrane permeabilization (MOMP)^3^. Initially discovered in chromosomal translocations in follicular lymphomas, Bcl-2 homologs have now been found in almost all metazoans. Indeed, genetic screening of cell death mutants performed in nematodes and subsequent studies in drosophila, zebrafish and hydra uncovered a number of Bcl-2-related proteins. ^4–7^. These proteins contain one to four B-cell homology domains, as well as a hydrophobic C-terminus transmembrane domain ^8^. Although the control of apoptosis differs substantially between roundworms, insects and mammals, seminal experiments conducted in these models reinforced the dogma that Bcl-2 family of proteins were evolutionary selected to control the survival of the cell at the level of the mitochondria. However, a number of Bcl-2-related proteins are localized not only to the MOM but are also found at the endoplasmic reticulum (ER). Indeed, Bcl-2 and related members bind to the ER membranes and control intracellular calcium fluxes through direct interactions with the calcium channel inositol 1,4,5-trisphosphate receptor (IP_3_R) ^9–11^. Calcium represents a key second messenger implicated in a broad range of processes including cell cycle, cell metabolism, cell migration and apoptosis, and Bcl-2 family members have also been linked to the control of some of these processes through the control of calcium fluxes ^12^.

Recently we showed that two Bcl-2 homologs named Nrz and Bcl-wav were able to orchestrate the first morphogenic movements, during early zebrafish embryonic development independently from their roles in apoptosis. Nrz exerts this non-canonical function at the ER by directly regulating IP_3_R permeability, whereas Bcl-wav binds to VDAC channel at the MOM and enhances mitochondrial calcium uptake ^11, 13, 14^. We demonstrated that these molecular interactions control actomyosin contractility and dynamics in the developing embryo. Thus, the assumption that Bcl-2 family proteins appeared and were selected throughout metazoan evolution as sole MOMP regulators is now challenged ^15^.

Metazoans appeared during the late Precambrian period around 780 million years ago ^16^. Placozoans represent one of the most basal branching metazoans with *Trichoplax adhaerens* being one of the three living representatives of this phylum ^17^. It possesses the simplest animal morphology and is composed of only six cell types ^18, 19^. It does not undergo embryonic development but rather divides by scission. Recently, the genome of *Trichoplax adhaerens* was sequenced ^20^.

Here we describe the identification and the molecular characterization of four *Trichoplax adhaerens* Bcl-2-like (trBcl-2L1 to -2L4) proteins. Phylogenetic analyses demonstrated that Trichoplax possesses two anti-apoptotic Bcl-2 homologs (trBcl-2L1 and -2L2) and two members clustering with the pro-apoptotic subgroup (trBcl-2L3 and -2L4). We demonstrated that trBcl-2L3 is a genuine Bax-like protein, whereas trBcl-2L4 is a Bak homolog. Interestingly, trBcl-2L3 is a pro-apoptotic protein capable of inducing MOMP in *bax/bak*-deficient baby mouse kidney (BMK) cells whereas trBcl-2L4 acts as BH3-only sensitizer repressing anti-apoptotic activities of trBcl-2L1 and trBcl-2L2.

Finally, we showed that trBcl-2L1 and trBcl-2L2 possess dual subcellular localizations at the ER and the mitochondria, and demonstrate that trBcl-2L1 controls IP_3_R-dependent calcium fluxes. Overall, these results suggest that Bcl-2 proteins may have emerged as multifunctional cell factors capable of controlling both MOMP and calcium fluxes.

## RESULTS

### In silico identification and characterization of Bcl-2-like proteins

We performed extensive tBlastN screening of the genome of the *Trichoplax adhaerens* Grell-BS-1999 strain publically available on the NCBI. Using the full length human Bcl-2 protein as well as individual BH and TM domains as a query we identified four protein encoding sequences potentially corresponding to Bcl-2 homologs. We named them *Trichoplax adhaerens* Bcl-2-like proteins or trBcl-2l (Figure 1A). We further confirmed the expression and the sequence of the four *trbcl2*-like genes in *Trichoplax adhaerens* by performing reverse transcriptase PCR using specific primers targeting the exon #1 of each gene (Figure S1A). Sequences were deposited into the NCBI databank under the following accession numbers KU500586.1, KU500587.1, KU500588.1, and KU500589.1 corresponding to the trBcl-2L1, trBcl-2L2, trBcl-2L3 and trBcl-2L4, respectively. Multiple alignments coupled with secondary structure prediction showed that all four sequences presented the typical multidomain BH organization, the TM domain located in the C-terminus end, as well as all alpha helices secondary structures (Figure 1A, Figure S1B). Interestingly, analysis of the primary structure of trBcl-2L3 also highlighted a unique S/T rich region between alpha helix 1 and 2. This region was predicted as a mucin-like domain which is found in secreted proteins. Of note, during the course of our survey we were not able to detect BH3-only encoding genes in the *T. adhaerens* genome.

**Figure 1.**
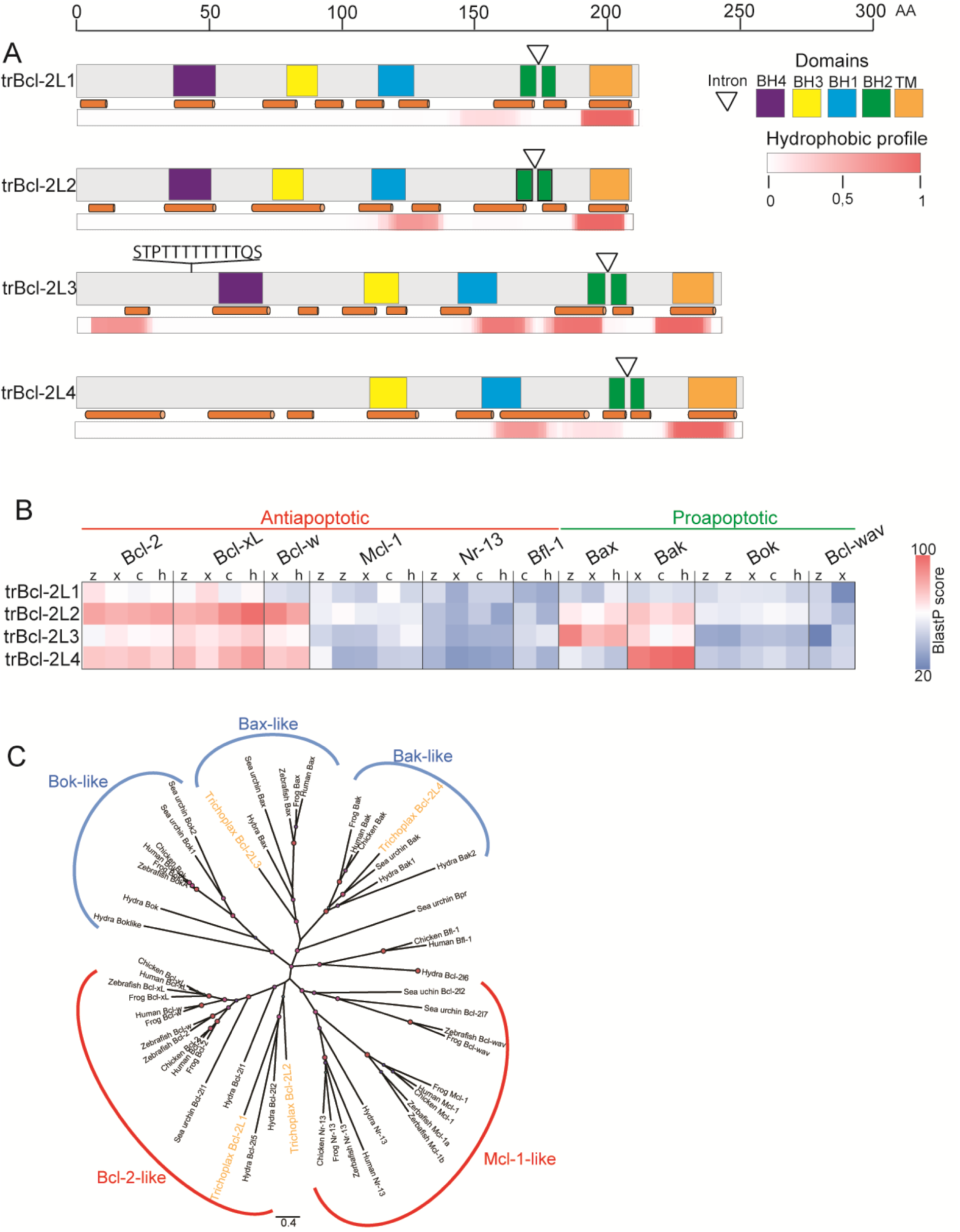
Sequence and expression analysis of Bcl-2 homologs found in *Trichoplax adhaerens*. **(A)** Schematic representation of the primary structures of four multidomain *Trichoplax adhaerens* Bcl-2-like (trBcl-2L1 to -2l4) proteins. The positions of conserved BH1-BH4 domains and of the C-terminus transmembrane domain (TM) as well as the locations of the predicted alpha helices are indicated with colored boxes and orange cylinders respectively. The white triangles represent the position of the conserved BH2 splitting intron. The hydrophobic profile for each protein is presented below. An amino acid scale bar is presented above. **(B)** *In silico* hybridization matrix representing the alignment of four trBcl-2-like proteins *versus* Bcl-2 related multidomain proteins found in *Danio rerio* (zebrafish = z), *Gallus gallus* (chicken = c), *Xenopus tropicalis* (xenopus = x), and *Homo sapiens* (human = h). Colored boxes refer to the total alignment score (blue; poor alignment, red; high alignment (> 60). trBcl-2L1 and trBcl-2L2 appear to cluster with anti-apoptotic members, whereas trBcl-2L3 and trBcl-2L4 with the pro-apoptotic Bax and Bak, respectively. **(C)** Phylogenetic tree using the neighbor joining clustering method. TrBcl-2L1 and trBcl-2L2 cluster with the Bcl-2-like subgroup whereas trBcl-2L3 and trBcl-2L4 cluster with Bax and Bak orthologs, respectively.

In effect, trBcl-2 homologs cluster in two groups at both genomic and protein levels. For instance all four trBcl-2-like genes possess a unique BH2 splitting intron. However this intron was relatively long for *trbcl2l1* and *trbcl2l2* with 1991 nt and 1725 nt, respectively whereas it was significantly shorter in *trbcl2l3* and *trbcl2l4* with 463 nt and 280 nt, respectively. Furthermore, trBcl-2L1 and trBcl-2L2 proteins had similar molecular weights and isoelectric points (pI): 24766 and 24282 Da and pI of 8.65 and 8.85, respectively, whereas trBcl-2L3 and trBcl-2L4 were slightly bigger, 27382 and 27944 Da with lower pI 6.96 and 6.45 respectively. Finally, using local alignment matrices and neighbor joining phylogenetic tree generation, we showed that trBcl-2L1 and trBcl-2L2 were able to cluster with the anti-apoptotic subgroup of the Bcl-2 family, whereas trBcl-2L3 and trBcl-2L4 clustered with the pro-apoptotic subgroup (Figure 1B&C). TrBcl-2L1 and -2L2 homologs were highly divergent, whereas trBcl-2L3 and trBcl-2L4 clustered well with Bax and Bak homologs, respectively. We hereafter named these proteins trBax and trBak, respectively.

### Role of trBcl-2 proteins in the control of MOMP

To gain further mechanical insight into the molecular activity of these trBcl-2-like proteins, we performed apoptosis assay in HeLa cells by analyzing the activation of Caspase 3 using a specific antibody. The transient expression of trBcl-2L1 (2.2% ± 1.3%) and trBcl-2L2 (2.4% ± 1.2%) for 24 h did not lead to a significant increase in Caspase 3 activation compared to control cells expressing the ER-targeted EGFP (6.3% ± 5.2). Conversely, the expression of trBax (89.5% ± 4.5) and trBak (88.7% ± 4.5) led a massive Caspase 3 activation (Figure 2A&S2A), close to that observed with the zebrafish orthologue of Bax (zBax) (87.0% ± 4.7), suggesting that these two Trichoplax proteins are genuine Bcl-2 pro-apoptotic members (Figure 2A&S2A). We further confirmed these results using zebrafish embryos as a model. Indeed, during zebrafish development, embryos are resistant to apoptotic stimuli prior to the gastrulation stage which starts approximately at 6 hours post fertilization (hpf). The forced expression of Bcl-2 pro-apoptotic proteins has been shown to induce apoptosis in these embryos prior to this stage ^7, 11^. We thus injected at the one cell stage *in vitro* transcribed mRNA encoding for each Bcl-2 homolog. Expression of trBcl-2L1 and -2L2 did increase embryo mortality. However, the expression of trBax and trBak led to embryo death, reaching almost 100% at 6 hpf (Figure 2B). Interestingly, co-expression of trBcl-2L1 and -2L2 in combination with the trBax and trBak proteins led to the complete inhibition of their pro-apoptotic activity. These results suggest that trBax and trBak are pro-apoptotic Bcl-2 members, the activities of which may be antagonized by trBcl-2L1 and -2L2 proteins probably through direct interactions. To test this hypothesis, we performed fluorescence resonance energy transfer (FRET)-based interaction analyses between an EGFP donor and an Alexa568 acceptor in BMK DKO cells. Indeed, we expressed EGFP-tagged trBax and trBak in combination with Flag-tagged trBcl-2L1 and trBcl-2L2 and performed immunofluorescence using anti-Flag and secondary antibodies coupled with Alexa568 fluorophore. Interactions between trBcl-2-like proteins were estimated by measuring the dequenching of EGFP emission induced by Alexa568 photobleaching. As seen on Figure 2C, FRET efficacy between EGFPtrBax/Flag-trBcl-2L1 (5.84%±0.84%) and EGFPtrBak/Flag-trBcl-2L2 (5.45%±0.65%) was similar to control cells expressing EGFP fused to human Bax and to Flag-tagged human Bcl-2 protein (4.92%±0.78%). Interestingly, FRET efficacy between EGFPtrBak/Flag-trBcl-2L1 (11.18%±1.34%) was close to the positive control a NeonGreen-Flag-Nrh (11.20%±0.93%) protein, whereas EGFPtrBax and Flag-trBcl-2L2 interacted only weakly (1.97%±0.96%) (Figure 2C&S2B).

**Figure 2.**
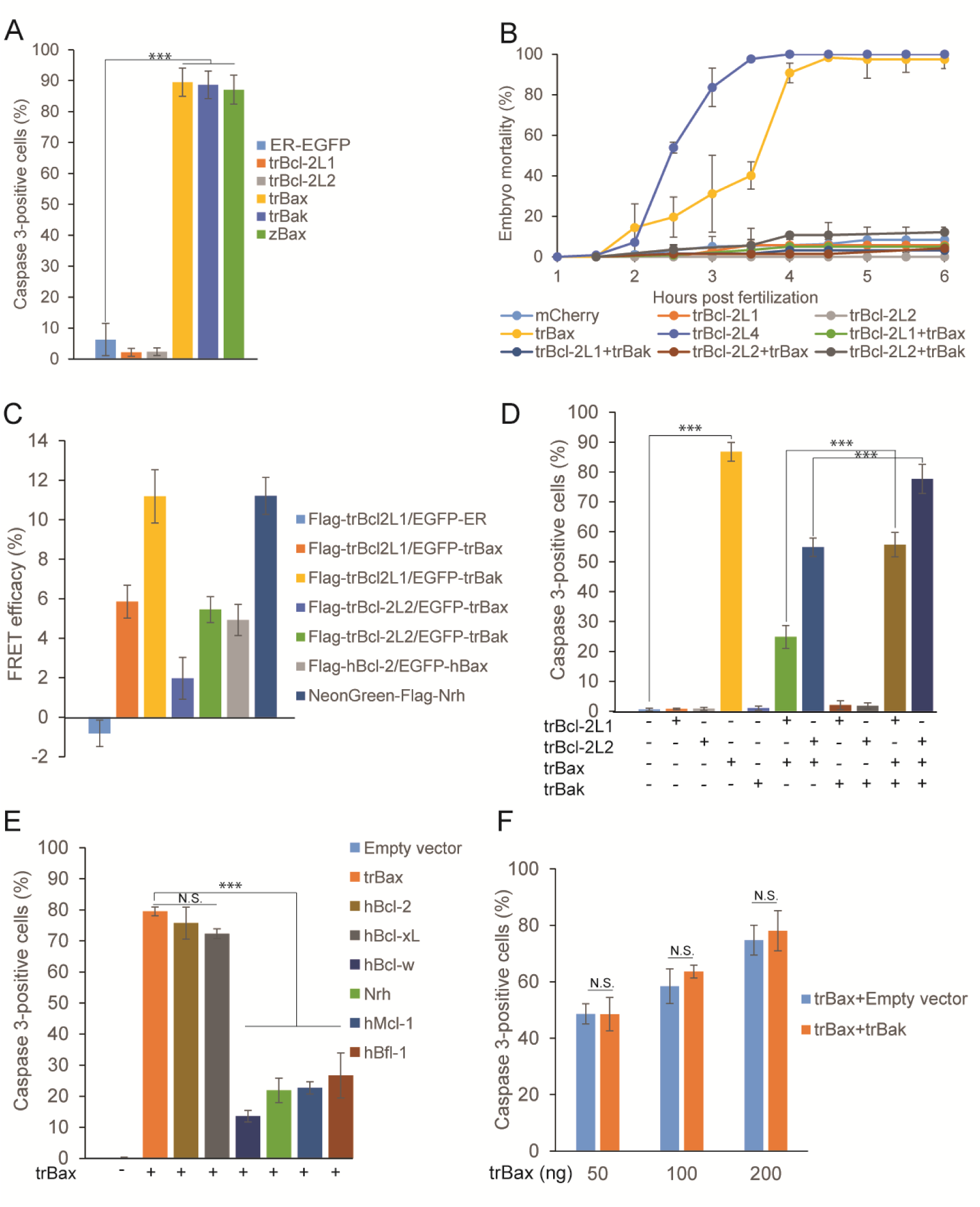
Control of MOMP by trBcl-2-like proteins. **(A)** Quantification of the Caspase 3 activation in HeLa cells transiently expressing trBcl-2-like proteins (mean ± SEM; three independent experiments). **(B)** Graph presenting the effect of the trBcl-2-like protein expression on the mortality of zebrafish embryos. In contrast to trBcl-2L1 and trBcl-2L2 which had no effect mRNA injection (100 ng/µL) into one cell stage embryos of trBax and trBak induced early mortality during gastrulation (mean ± SEM; three independent experiments). **(C)** Graph showing the results from FRET-based interaction analyses between an EGFP donor and an Alexa568 acceptor. BMK DKO cells expressing EGFP-tagged trBax and trBak in combination with Flag-tagged trBcl-2L1 and trBcl-2L2 were fixed and labeled using anti-Flag and secondary anti-IgG mouse antibodies coupled with Alexa568. Interactions between trBcl-2-like proteins were estimated by measuring the dequenching of EGFP emission induced by Alexa568 photobleaching. **(D)** Histogram showing the effect of trBcl-2-like protein expression on Caspase 3 activation in BMK double *bax -/-, bak-/-* knock out (DKO) cells. In comparison to HeLa cells, trBak protein is unable to induce Caspase 3 activation in BMK DKO cells (mean ± SEM; three independent experiments). **(E)** Histogram showing the effect of the expression of human Bcl-2 homologs on the pro-apoptotic activity of trBax protein in BMK DKO cells. Proteins belonging to the Mcl-1 subgroup as well as Bcl-w antagonize trBax pro-apoptotic activity, while Bcl-2 and Bcl-xL have no effect. (mean ± SEM; three independent experiments). **(F)** Histogram showing the effect of the expression of trBak protein on the pro-apoptotic activity of trBax in BMK DKO cells. Cells were transfected with increasing amounts of trBax expressing vector 50, 100 and 200 ng in the presence of constant amount of pCS2+ empty vector or pCS2+trBak. TrBak coexpression did not lead to increase of the percentage of Caspase 3-positive cells compared to trBax expressing vector plus empty vector (mean ± SEM; three independent experiments)

We next evaluated the capacity of trBax to induce apoptosis independently of the presence of endogenous Bax and Bak by performing a series of apoptosis assays using baby mouse kidney cells knocked out for endogenous *bax* and *bak* (BMK DKO cells) (Figure S2C). When expressed in these cells, EGFP tagged trBax rapidly translocated to the mitochondria where it induced mitochondrial fragmentation, Δψ loss and Cytochrome C release (Figures S2D-H). Furthermore the majority of the trBax expressing cells were Caspase 3-positive (86.8% ± 3.1%) in comparison to control cells (0.5% ± 0.45%), demonstrating that trBax is a *bona fide* Bax-like protein (Figure 2D). Of note, trBcl-2L1 (24.8% ± 3.8%) and to lesser extent trBcl-2L2 (54.9% ± 3.1%) were able to counteract the pro-apoptotic activity of trBax (Figure 2D).

However, inhibition of trBax pro-apoptotic activity was not restricted to Trichoplax Bcl-2 members. Indeed, when human Bcl-2 homologs were expressed in combination with trBax we observed a significant decrease in the number of Caspase 3-positive cells (Figure 2E&S2I). This was however not the case for all human anti-apoptotic Bcl-2 proteins. For instance Mcl-1, Bfl-1, Nrh and Bcl-w inhibited trBax in a similar manner as trBcl-2L1, while Bcl-2 and Bcl-xL had no effect on its pro-apoptotic activity, suggesting an evolutionary conservation in the interaction between the hydrophobic BH3 domain-binding groove of anti-apoptotic Bcl-2 proteins and the BH3 domain of trBax.

Next, we evaluated the effect of trBak expression on BMK DKO apoptotic cell death. Although trBak was highly expressed in BMK DKO cells, it did not result in Cytochrome C release or significant Caspase 3 activation showing that the pro-apoptotic activity of trBak is dependent on the presence of endogenous Bax or Bak (Figure 2D&S2C-E, S2G). Interestingly, trBak co-expression inhibited the anti-apoptotic activity of trBcl-2L1 and trBcl-2L2 over trBax, without increasing its pro-apoptotic potential suggesting that this protein actually acts as a BH3-only sensitizer (Figure 2D&2F).

### Subcellular localization of trBcl-2 proteins

In vertebrates, members of the Bcl-2 family can be localized both at the level of the mitochondria and ER. In order to test whether this feature was also present in Trichoplax Bcl-2 homologs, we transiently expressed in mammalian cells, N-terminus Flag-tagged trBcl-2-like proteins in combination with ER-targeted EGFP. Under these conditions we were able to detect trBcl-2L1 and trBcl-2L2 in mitochondrial and ER membranes (Figure 3A). Of note cells expressing trBcl-2L2 proteins presented hyper fused perinuclear mitochondria which presented a decreased membrane potential (data not shown). In addition, intracellular trBcl-2L2 localization was more dispersed than trBcl-2L1, suggesting that this protein could be also cytosolic. These observations were confirmed by performing subcellular fractionation on BMK DKO cells followed by immunoblotting. Indeed trBcl-2L1 was found to be membrane bound whereas a fraction of trBcl-2L2 was found in the cytosol (Figure 3B). When we expressed trBax and trBak we detected these proteins majorly at the level of the mitochondria with minor expression in the microsomal fraction (Figures 3A&B).

**Figure 3.**
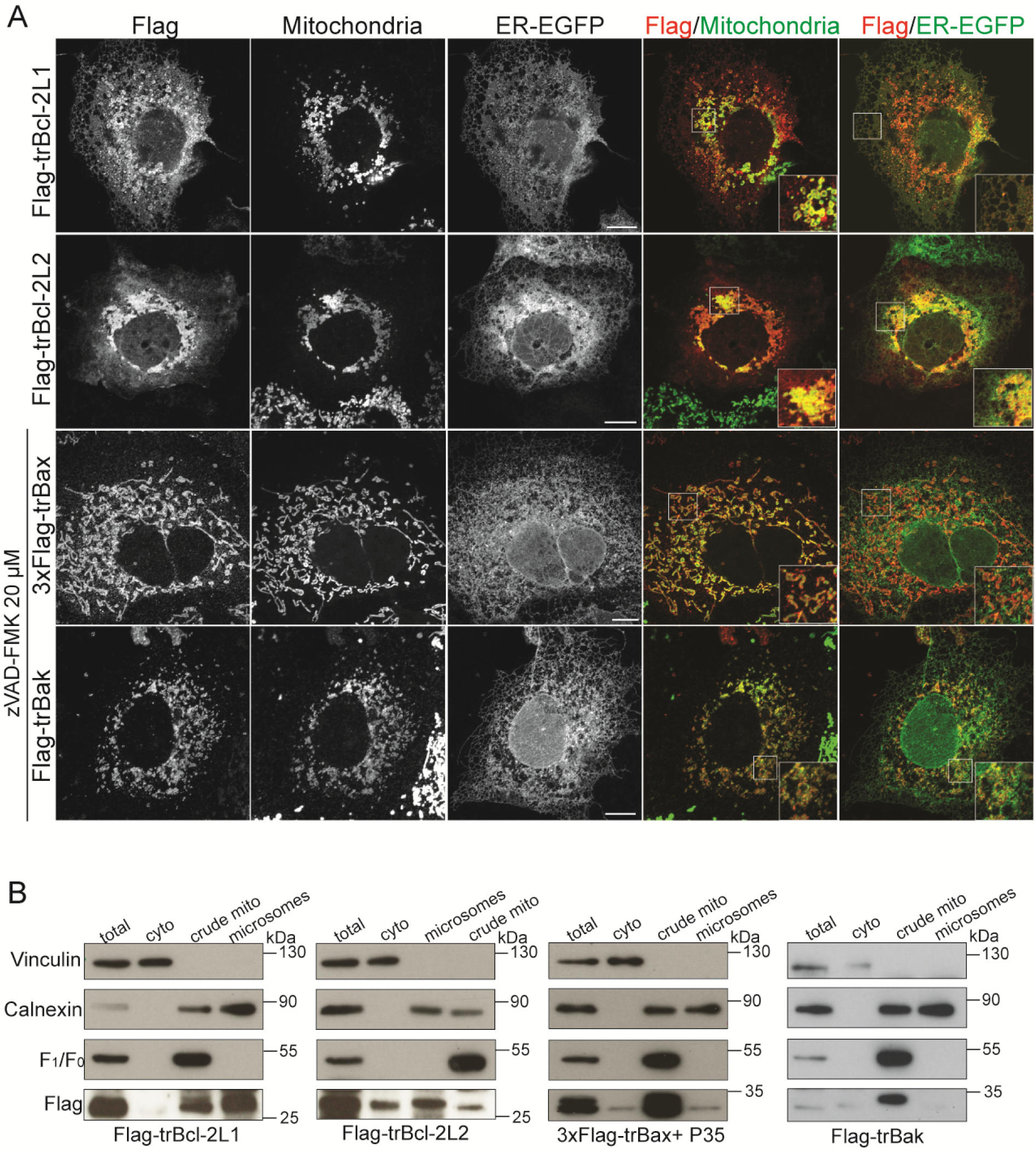
Subcellular localization of trBcl-2-like proteins. **(A)** Representative confocal images of Cos-7 cells expressing trBcl-2-like proteins. Subcellular localization of each protein was analyzed using ER-targeted EGFP for staining ER membranes, the Mitotracker™ deep red dye for labeling mitochondria, and the anti-Flag antibody for the detection of Flag tagged Bcl-2-like proteins. Merged channels between Flag and ER-GFP and between Flag and Mitochondria were presented on the right. White boxes represent the enlarged area of each merged image. Scale bars: 20 µm. **(B)** Subcellular fractionation of BMK DKO cells expressing Flag-tagged trBcl-2-like proteins. TrBcl-2L1 and trBcl-2L2 proteins were found in crude mitochondrial and microsomal fractions. Full length trBcl-2L2 was also detected in the cytosol fraction. TrBax and trBak were found mostly in the crude mitochondria fraction. Anti F_1_F_0_ ATPase, Calnexin and Vinculin antibodies were used as mitochondrial, ER and cytosol markers, respectively.

We next performed experiments in in the budding yeast *Saccharomyces cerevisiae*. Treatment of purified yeast mitochondria with Na_2_CO_3_ showed that trBcl-2L1 is tightly inserted in the mitochondrial membrane whereas trBcl-2L2 and trBak are more loosely attached (Figure S3). Interestingly, co-expression with P168A mutant of human Bax, which is integral mitochondrial protein leads to trBcl-2L2 and trBak stabilization at the MOM. Of note expression of trBax was highly cytotoxic and did not allowed the detection of the protein.

Bax localization to MOM represents a dynamic process of translocation and retrotranslocation which depends on internal cues most importantly the α9 hydrophobic helix containing the TM domain but also the α1 helix and a proline residue located after the BH2 domain ^21–23^. It has been shown that the point mutation of this residue at position 168 into alanine actually impairs the proper translocation-retrotranslocation dynamics of human Bax (hBax) ^23, 24^. Interestingly, trBax WT is a natural alanine mutant (alanine at position 212) which is constantly located at the MOM (Figures S2F&H). It possesses α1 and α9 hydrophobic helices as well as a unique S/T rich region between α helix 1 and 2, predicted as a mucin-like domain, found in secreted proteins. In order to understand how these residues regulates trBax localization at the MOM we generated two site-directed mutants: P212A and N(8)T in which we replaced the alanine residue with proline and the eight threonine residues by asparagine residues, respectively. We also generated three trBax deletion mutants, lacking the α1 (Δα1), α9 (Δα9) or both α1and α9 (Δα1Δα9) helices (Figure S4A). Expression of EGFP-tagged WT trBax rapidly resulted in its translocation to the mitochondria in comparison with hBax in which a significant fraction remained cytosolic suggesting that trBax does not possess the same translocation-retrotranslocation dynamics as human Bax does (Figure S4B, Figures S2F&H).

When we expressed trBax P212A and N(8)T mutants we did not detect any changes in the subcellular distribution compared to WT trBax. However mutants lacking α9 helix but not α1 had clear cytosolic localization. Of note the mitochondrial localization of trBax was tightly correlated with its capacity to induce Caspase 3 activation as mutants lacking the TM domain had reduced capacities to induce cell death in BMK DKO cells (Figure S4C).

The importance of the TM domain for the subcellular localization of the other trBcl-2 proteins was further demonstrated. To this end we generated mutants lacking the last hydrophobic alpha helix (ΔTM) predicted *in silico*. When expressed in Cos-7 cells the trBcl-2L1ΔTM, L2ΔTM and trBakΔTM mainly displayed a cytosolic distribution (Figure S4D).

### TrBcl-2L1 has a dual activity at the mitochondrion and ER

Since two members of the Bcl-2 family in Trichoplax have a dual ER/mitochondria subcellular localization, we hypothesized that these proteins may control intracellular calcium fluxes at the level of the ER. We focused on trBcl-2L1 which possesses a clear ER and mitochondrial localization without proteolytic cleavage. To test the activity of trBcl-2L1 at the ER we expressed this protein in combination with the ER targeted Ca^2+^ biosensor R-CEPIA1er and subject the cells to IP_3_-dependent calcium release using histamine. In comparison with control cells (0.60 ± 0.051) which were transfected with the empty vector, trBcl-2L1-expressing cells had a lower level of intraluminal calcium (0.37 ± 0.082) following histamine treatment, suggesting that trBcl-2L1 is able to increase the permeability of the IP_3_R calcium channel (Figure 4A-C). This calcium release from the ER led to concomitant increase in cytosolic (1.58 ± 0.02) and mitochondrial matrix Ca^2+^ levels (1.60 ± 0.02) compared to control cells (1.35 ± 0.04) (1.47 ± 0.034) (Figure S5A-D). In order to test whether trBcl-2L1 is able to bind to IP_3_R we performed a co-immunoprecipitation assay in HeLa cells transiently transfected with Flag-trBcl-2L1. As shown on Figure 4D trBcl-2L1 co-immunoprecipitated with endogenous IP_3_R type 1 supporting that the calcium regulation is indeed IP_3_R dependent.

**Figure 4.**
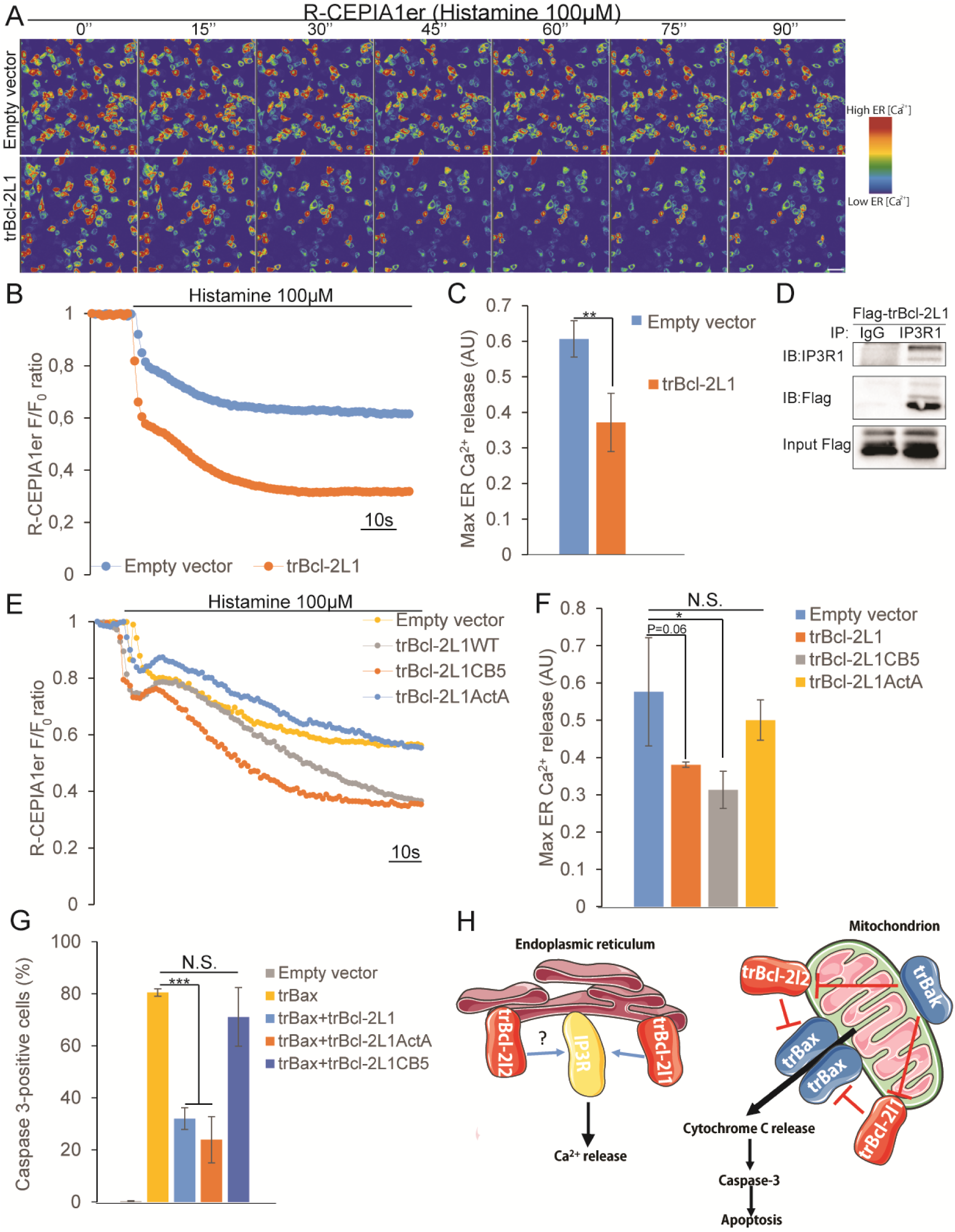
Dual role of the trBcl-2L1 protein at the ER and mitochondrion. **(A)** Representative images of HeLa cells expressing ER targeted R-CEPIA1er biosensor in combination with pCS2+ empty vector or pCS2+ vector encoding trBcl-2L1. Cells were treated with histamine (100 µM) to induce Ca^2+^ depletion in the ER. **(B)** Traces showing the relative changes in R-CEPIA1er fluorescence intensities normalized against the baseline value (F/F_0_ ratio). Histamine (100 µM) stimulation induces ER Ca^2+^ depletion. Compared to control cells, trBcl-2L1-expressing cells possess higher ER releasing capacities. **(C)** Histogram depicting the maximum ER release (Fmax-Fmin difference) of HeLa cells expressing trBcl-2L1 or transfected with empty vector. (mean±S.E.M; n = 3 independent experiments). ** *P* < 0.01. **(D)** Immunoprecipitation experiments with protein extracts from transfected HeLa cells with Flag-trBcl-2L1 co-immunoprecipitated with the IP3R1 channel **(E)** Representative curves showing the relative changes in R-CEPIA1er fluorescence intensities normalized against baseline values (F/F_0_ ratio) following histamine (100 µM) treatment. ER-targeted trBcl-2L1 and WT trBcl-2L1 have similar releasing capacities, whereas Mito-targeted trBcl-2L1 resembles cells transfected with empty vector. **(E)** Histogram depicting the maximum ER release (Fmax-Fmin difference) of HeLa cells expressing WT trBcl-2L1 or single localization mutants (mean ± S.E.M; n = 3 independent experiments). * *P* < 0.05. N.S. not significant. **(F)** Quantification of the Caspase 3 activation in HeLa cells expressing trBax alone or in combination with trBcl-2L1 WT protein or dedicated single localization mutants (mean ± S.E.M; n = 3 independent experiments). *** *P* < 0.001. N.S. not significant. (mean ± SEM; three independent experiments). **(G)** Schematic representation of the model of action of trBcl-2 like proteins. Anti-apoptotic trBcl-2L1 and L2 proteins are able to locate to the mitochondria and the ER. At the level of the mitochondria trBcl-2L1 and L2 bind and inhibit trBax pro-apoptotic activity whereas trBak acts as repressor of their anti-apoptotic activity allowing trBax monomers to accumulate and form pores at the MOM. Independently of this function, trBcl-2L1 is also able to attach to the ER membrane where it controls IP3R dependent calcium release.

We next generated two single subcellular mutants by replacing its endogenous TM domain with the TM domain of the ER-based Cytochrome B5 protein or with the TM domain of the MOM-localized ActA protein (Figure S5E-H). Expression of trBcl-2L1CB5 but not trBcl-2L1ActA was able to sustain higher IP3R dependent calcium release demonstrating that indeed this molecular function of trBcl-2L1 depends on its ER-localization (Figure 4E&F). Interestingly, when we tested the capacities of each trBcl-2L1 mutant to repress the pro-apoptotic activity of trBax we observed the opposite result (Figure 4G); TrBcl-2L1ActaA was able to repress trBax-induced apoptosis, whereas trBcl-2L1CB5 had no effect.

Altogether these results suggest that at the molecular level trBcl-2 proteins possess distinct functions in the control of the MOMP or in the regulation of intracellular calcium fluxes which are predetermined by their subcellular localization (Figure 4H).

## DISCUSSION

During the course of our research we identified and molecularly characterized four Bcl-2 homologs encoded by the genome of *Trichoplax adhaerens*. All four genes harbor a single conserved BH2 splitting intron, which is located specifically after the TGG codon encoding a tryptophan residue. This specific intron is not restricted to placozoans but has been found in almost all *bcl2* homologous genes in multicellular animals ^25^. This observation suggests that Bcl-2 multidomain members arose from a common *bcl2* ancestral gene which already possessed this BH2 splitting intron. However, it remains unclear whether this ancestral gene had pro- and/or anti-apoptotic functions or even functions unrelated to the regulation of MOMP. Mining the Trichoplax genome did not allow us to detect a gene encoding a BH3-only protein. One possibility is that Trichoplax encodes for a highly divergent BH3-only protein which currently remains elusive. Alternatively, it is possible that trBak, although structurally related to multidomain Bcl-2 proteins, plays the role of a functional BH3-only protein. Indeed, when expressed in BMK DKO cells, trBak failed to induce Cytochrome C release and Caspase 3 activation in comparison with trBax which promoted MOMP independently of the endogenous Bax and Bak status. Furthermore, trBak binds to both anti-apoptotic trBcl-2L1 and -2L2. This suggests that trBak protein may function as a BH3 only sensitizer that repress heterodimers formation between anti-apoptotic trBcl-2 proteins and trBax which ultimately results in trBax oligomerization. Indeed, our results show that trBax is a *bona fide* pro-apoptotic Bax like protein that is majorly localized to the MOM, where induces MOMP and cytochrome C release in cells lacking endogenous Bax and Bak. Interestingly, unlike its human ortholog which dynamically navigates between the cytosol and the MOM though translocation and retrotranslocation, trBax seems to directly target the mitochondrion. This mitochondrial localization depends on the C-terminus TM domain. Indeed, trBax mutants lacking this hydrophobic helix mainly displayed a cytosolic distribution and a decreased pro-apoptotic activity. Interestingly, deletion of both α1 and α9 helices completely abolished the capacity of trBax to induce Caspase 3 activation, suggesting that the hydrophobic α1 helix may also contribute to trBax localization to the MOM.

TrBax protein also possesses a serine/threonine rich sequence located between the α1 and α2 helices. I*n silico* prediction identified this region as a mucin-like domain. Mucins are secreted proteins subjected to O-linked glycosylation at these serine/threonine residues at the level of the Golgi apparatus. Although mutation of these residues into asparagine residues did not affect trBax subcellular localization, it is tempting to speculate that *in vivo* trBax might function as an excreted cellular toxin. Thus it would be of interest to detect the subcellular localization of trBax and determine its glycosylation profile in Trichoplax.

Our current results strongly suggest that in Trichoplax, trBcl-2 proteins may contribute to control apoptosis through the regulation of MOMP. Indeed, analysis of the Trichoplax genome revealed the presence of key components of the mitochondrial pathway including Apaf-1, Cytochrome C as well as initiator and effector caspases ^20, 26^. However, whether apoptosis occurs in Trichoplax *in vivo* remains unclear. Research performed on this model using nutlin-3 a cis-imidazoline analog which inhibits the interaction between Mdm2 and p53 or roscovitine a cyclin-dependent kinase inhibitor were shown to induce TUNEL-positive cells, indicating that at least in stress conditions apoptosis may operate in placozoans ^27^. Placozoans are small animals that lack embryonic development and reproduce mainly vegetatively though fission or budding. They are commonly found in coastal zones in tropical and subtropical waters ^28^. Thus, it is tempting to speculate that these animals may commonly be exposed to various stresses including UV radiation, or mechanical and chemical stresses, suggesting that apoptosis may play an important role in the maintenance of homeostasis.

We finally demonstrated that the anti-apoptotic Trichoplax Bcl-2 members possess double ER and mitochondrial subcellular localization, indicating that they may act at the level of the ER especially in the control of the intracellular calcium levels. This was actually the case for trBcl-2L1, which is able to increase the permeability or IP_3_R stimulated by histamine. Interestingly, this function required the sole ER localization of trBcl-2L1, whereas the mitochondrial localization is required for inhibiting trBax MOMP. These results demonstrate that at the molecular level members of the Bcl-2 family are actually multifunctional and that the interaction between Bcl-2 family proteins and calcium related channels such as IP3R appeared early in the evolution of metazoans. Future research should be performed on Trichoplax animals in order to analyze the implications of these two functions *in vivo*.

## MATERIALS AND METHODS

### TrBcl-2 ORF cloning and expression analysis

Approximately 100 randomly picked animals of *Trichoplax adhaerens* “Grell” were used for total RNA isolation (for animal culture conditions, see ^29^). The animals were initially transferred into a clean glass petri dish, washed three times with 3.5% artificial seawater (ASW) and afterwards starved overnight to prevent algae contamination. The animals were transferred into a 1,5ml Eppendorf tube and the ASW was removed after short centrifugation. Animals were lysed in 500µl pre-warmed homogenization buffer (HOM-I: 100mM Tris HCl, 10mM EDTA, 100mM NaCl, 2,5mM DTT, 0,5% SDS, 0.1% DEPC in ultra pure water at pH 8.0 and 65°C). Proteins were digested with 25µl DEPC-treated Proteinase K (10mg/ml) for 20 minutes at 65°C. The lysate was further homogenized by 2x suction through a hypodermic needle connected to a 2.5ml syringe. Nucleic acids were isolated from the lysate by a standard Phenol/Chloroform/Isoamylalcohol (25:24:1) extraction and a subsequent precipitation with Isopropanol. The following DNA digestion with DNase I (Thermo Fisher Scientific) was conducted according to the manufacturer’s recommendations. The quality and purity of total RNA was afterwards checked on an agarose gel. First-strand cDNA synthesis with oligo(dT) primers was conducted using the SuperScript II Reverse Transcriptase (Thermo Fisher Scientific) following the manufacturer’s protocol.

The open reading frames of the four Bcl-2 homologs in Trichoplax were cloned using cDNA from the *Trichoplax adhaerens* Grell clone (kindly provided from Bernd Schierwater lab) using the following primers:

trBcl-2L1-F 5’ATGTCTCCTGAACGCGAAAAAATGG

trBcl-2L1-R 5’TCACCATCTTTCCCACACTGCAAGCG

trBcl-2L2-F 5’ATGACATCTTCCTTCGATGAGGCATC

trBcl-2L2-R 5’TTATCTATTAAACAGAAAGTATGCACAAGTACC

trBcl-2L3-F 5’ATGGCCGATTTCACGTATGTACTGA

trBcl-2L3-R 5’TTAGCTACTGTTATATCGTTGCCATGTG

trBcl-2L4-F 5’ATGTCTAGCACTATTACAATAGCTGAATCGC

trBcl-2L4-R 5’TTATTCTCTAAAATAGCCTGAAAGCCATAGG

The PCR amplicons obtained were cloned into the pJET1.2/blunt cloning vector (Thermo Fisher Scientific) and sent for sequencing. Verified sequences were deposited in an NCBI databank with the following Acc. Numbers KU500586.1 (trBcl-2L1), KU500587.1 (trBcl-2L2), KU500588.1 (trBcl-2L3), KU500589.1 (trBcl-2L4).

For expression analysis internal primers targeting exon 1 of each *bcl2* homolog were used for PCR amplification with the following sequences:

Int-trBcl-2L1-F 5’TCCGTAATGGCTGCAACTGG

Int-trBcl-2L1-R 5’CTCACGTACTCATCCCAGCC

Int-trBcl-2L2-F 5’GCTTACCCAACGTTTGTGGG

Int-trBcl-2L2-R 5’TGTTCGCTTCGATCCAGGTAG

Int-trBcl-2L3-F 5’CCACCACTACTCAGTCATGTCC

Int-trBcl-2L3-R 5’TGTTGGTGACGTCGGTTCAA

Int-trBcl-2L4-F 5’CTATCGGCAAATCCGGCTCA

Int-trBcl-2L4-R 5’TGCGCTAAATGGCGTCCTAT

Flag-tagged sequences were ordered using G-blocks synthesis (IDTDNA) and were cloned into the pCS2+ expression vector between BamHI/XhoI. TrBax sequence was codon optimized for expression in eukaryote cell system. These sequences were used for PCR amplification. HA-tagged trBcl-2-like and 3xFlag-tagged sequences were cloned into the pCS2+ expression vector between BamHI/XhoI and EcoRI/XhoI restriction sites respectively. TrBak, trBax and associated mutants were cloned in pEGFP-C1 vector using BglII/EcoRI restriction sites using the following primers:

HA-trBcl-2L1-F

5’ATATGGATCCATGTACCCATACGATGTTCCAGATTACGCTATGTCTCCTGAACGCGAAAAAATGG

HA-trBcl-2L1-R

5’ATATCTCGAGTCACCATCTTTCCCACACTGCAA

HA-trBcl-2L2-F

5’ATATGGATCCATGTACCCATACGATGTTCCAGATTACGCTATGACATCTTCCTTCGATGAGGCATC

HA-trBcl-2L2-R

5’ATATCTCGAGTTATCTATTAAACAGAAAGTATGCACAAGTACC

HA-trBax-F

5’ATATGGATCCATGTACCCATACGATGTTCCAGATTACGCTATGGCCGATTTCACGTATGTACTGA

HA-trBax-R

5’ATATCTCGAGTTAGCTACTGTTATATCGTTGCCATGTG

HA-trBak-F

5’ATATGGATCCATGTACCCATACGATGTTCCAGATTACGCTATGTCTAGCACTATTACAATAGC

HA-trBak-R

5’ATATCTCGAGTTATTCTCTAAAATAGCCTGAAAGCCATAGG

3xFlag-trBax-F

ATATGAATTCGCCGATTTCACGTATGTACTGA

3xFlag-trBax-R

ATATCTCGAGTTAGCTACTGTTATATCG

EGFP-deltaα1-trBaxopt-F

GTGTAGATCTAAGTCCGTGACTTCCACACCAACGA

EGFP-trBaxopt-R

ATATGAATTCCTAGCTGCTGTTGTAGCGCTGC

EGFP-deltaTM-trBaxopt-R

ATATGAATTCCTAGCTGCTGTTGTAGCGCTGC

P212A-F

TATATACAGCAACCGGACTGGAAGAACAAAT

P212-R

ATTTGTTCTTCCAGTCCGGTTGCTGTATATA

### Cell lines and transfection

HeLa, Cos-7, BMK WT or DKO cell lines were grown under standard cell culture conditions (37°C, 5% CO_2_) in Dulbecco’s modified Eagle’s medium (DMEM) high glucose medium (Gibco) supplemented with 10% FBS (Sigma), 100 U/mL penicillin and 100 µg/mL streptomycin (Gibco). Cell transfection was performed using X-tremeGENE™ HP DNA Transfection Reagent (Roche) according to the manufacturer’s protocol. Six hours post transfection cell medium was removed and cells were incubated for 24-48 h with new medium.

### Subcellular fractionation

All steps were carried out at 4°C. Transfected BMK DKO cells placed in two 15 cm in diameter plates were washed twice in PBS and added to 1,5 mL of ice cold MB buffer (210mM mannitol, 70 mM sucrose, 1 mM EDTA and 10 mM HEPES (pH 7.5) containing protease inhibitors). Cells were disrupted by shearing with a dounce homogenizer for 20 to 30 times. The disrupted cells were then centrifuged twice at 600 g for 5min to eliminate cell nuclei and centrifuged at 7,000 g to pelled crude mitochondria. Pellet was washed two times with fresh MB buffer and centrifuged at 10,000 g. Mitochondria were resuspended in an appropriate volume of MB buffer for further analyses. The supernatant was centrifuged at 100,000 g for 1 h, and the pellet containing the microsomal fraction was resuspended in RIPA buffer (150 mM NaCl, 1% Nonidet P-40, 0.5% Sodium deoxycholate, 0,1% SDS, 50 mM Tris-HCl (pH 7.4), containing protease inhibitors). Fifteen µg of each subcellular fraction and 50 µg of the total fraction were analyzed by western blot.

### Western blot analysis and antibodies

Western blots were performed as previously described ^30^. The following antibodies were used: F_1_F_0_ ATPase (BD #612518; dilution: 1/1,000), Vinculin (Santa Cruz #sc-55465; dilution: 1/2000), Calnexin (Cell Signaling Technology # 2679; dilution: 1/1000), Flag M2 (Sigma # F3165; dilution: 1/1,000), HA (Ozyme # BLE901501; dilution: 1/1,000), Cytochrome C (BD Bioscience # 556433; dilution: 1/1000). HRP-conjugated goat anti-mouse, rabbit anti-mouse and goat anti-rabbit secondary antibodies (DAKO) were used as secondary antibodies. Western blot analysis was performed according to standard procedures.

### Immunofluorescence staining, FRET measurements, cytochrome C release and apoptosis detection

Cells were placed on a glass coverslip in 12-well plates at 50% confluence and left to attach overnight. Cells were then transfected with trBcl-2-like encoding vectors in combination or pCS2+EGFP-CytB5 (ER-localized EGFP) for ER staining. For trBax and trBak subcellular localization zVAD-FMK was added 6 h post-transfection in order to delay cell detachment. For direct mitochondrial visualization cells were incubated with Mitotracker™ deep red dye (1/10,000) 20 min at 37°C. Cells were then washed once with medium for 20 min and then fixed with a 4% paraformaldehyde solution for 20 min followed by 3 washes with PBS. Following cell permeabilizaion using PBS Triton X-100 0.1% (PBT) plus BSA 3%, cells were incubated with primary antibodies Flag M2 (Sigma # F3165; dilution: 1/2,500), Cytochrome C (BD Bioscience # 556433; dilution: 1/1000) or Tom20 (Santa Cruz # sc-11415; dilution: 1/500). Goat anti-mouse or goat anti-rabbit IgG Alexa Fluor 488 (green fluorescent) or 568 (red fluorescent) were used as secondary antibodies (Molecular Probes) and used at 1/1,000 dilution for 1 h. Nuclei were visualized using Hoechst 33342 dye (Invitrogen #H3570) at 10 µg/mL. For FRET-based interaction analyses BMK DKO cells were transfected with Flag-tagged trBcl-2L1 and L2 in combination with EGFP-tagged trBax and trBak. Twenty four hours later cells were fixed and immunostained using mouse anti-Flag M2 antibody coupled anti-mouse Alexa568 secondary antibody. FRET was measured as previously described ^9^ Briefly EGFP fluorescent intensities were measured before and after photobleaching Alexa568 fluophore. FRET efficacy was calculated using the following formula 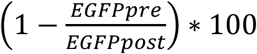

For apoptosis detection HeLa and BMK DKO cells transiently transfected with trBcl-2like proteins for 24 hours. Cells were fixed and permeabilized as stated above and incubated with anti-activated Caspase 3 antibody (BD Bioscience # 559565; dilution: 1/500). Pyknotic nuclei were labeled using Hoechst 33342 dye. Cells were observed under a fluorescent microscope and the percentage of Caspase 3- and Hoechst-positive cells was determined.

### Ca^2+^ measurements

ER calcium releasing capacities in BMK DKO stably expressing trBcl-2L1 or empty vector were measured using the ER probe R-CEPIA1er recombinant protein. Cells were transfected with pCMV R-CEPIA1er (Addgene #58216). Twenty-four hours later, cells were seeded onto 96-well plates and incubated overnight. Cells were washed with Ca^2+^-free balanced salt solution (BSS) [121 mM NaCl, 5.4 mM KCl, 0.8 mM MgCl_2_, 6 mM NaHCO_3_, 5.5 mM D-Glucose, 25 mM Hepes (pH 7.3)] and incubated with 180 µL of BSS buffer. Raw fluorescence values were collected every 0.21 sec for 2 min at 28°C using a Clariostar microplate fluorescence reader (BMG Labtech). After 5 sec of basal line measurement, 100 µM histamine (Sigma) was injected.

Cytosolic Ca^2+^ levels were detected in HeLa cells using Fluoforte dye (Enzo Life Sciences). Cells seeded in 96-well plates were incubated with 5 µM Fluoforte for 1 hr at 37°C, in a Ca^2+^-free BSS. Raw fluorescence values were collected every 0.21 sec for 2 min at 28°C using the Clariostar microplate reader. Mitochondrial calcium measurements were performed using Rhod-2 AM chemical dye or CEPIA2mt recombinant protein. Rhod-2 detection was performed as previously described for Fluoforte with the following modifications. Rhod-2 AM dye was incubated at 2.5 µM with 0.2% of pluronic acid (Molecular probes) for 20 min at 37°C in BBS-Ca^2+^ (BBS supplemented with 2 mM of Ca^2+^), and washed for 20 min with BBS-Ca^2+^ at room temperature. Raw fluorescence values were collected every 0.21 sec for 2 min at room temperature using the Clariostar microplate reader.

For CEPIA2mt detection, BMK DKO cells were co-transfected with pCS2+Flag-trBcl-2L4 and pCMV CEPIA2mt (Addgene #58218) and seeded at 40% in 8 well Nunc™ Lab-Tek™ II Chamber Slide. Twenty-four hours later, cells were washed with BBS-Ca2+. After 10 sec of baseline, 100 µM of histamine was added. Raw fluorescence values were collected every 0.21 sec for 2 min at room temperature using the Clariostar microplate reader.

### Zebrafish experiments

Zebrafish (crosses between AB and TU strains) were raised and maintained at 28.5°C according to standard procedures ^31^. Embryos were collected after fertilization and injected at the one-cell stage with *in vitro* transcribed mRNA encoding trBcl-2L1 to -2L4 proteins (100 pg total mRNA/ embryo) using mMESSAGE mMACHINE SP6 Transcription Kit (Thermo Fischer). Embryo death was monitored at regular intervals for 6 h post-fertilization.

### Yeast experiments

The 4 Trichoplax cDNAs encoding proteins having a Flag tag at their N-terminal end were cloned between the BamHI/XhoI sites of the pYES2 plasmid downstream to the inducible promoter GAL1/10.

The diploid yeast strain W303 (mat a/mat α, ade1/ade1, his3/his3, leu2/leu2, trp1/trp1, ura3/ura3) was used in all the experiment. Cells were co-transformed with pYES2-TrBcl-2like plasmids and pYES3-Bax plasmids. Two Bax variants were used: BaxWT coding the full-length untagged human protein, and BaxP168A carrying the P168>A substitution that converts Bax into a constitutively membrane-inserted and active protein ^24, 32^.

Transformants were selected on YNB plates supplemented with glucose and without uracil and tryptophan. For Bax and trBcl-2like proteins expressions, cells were grown aerobically at 28°C in YNB liquid medium (Yeast Nitrogen Base, 0.17%, potassium phosphate 0.1%, ammonium sulfate 0.5%, Drop-Mix 0.2%, auxotrophic requirements 0.01%) supplemented with 2% DL-lactate as a carbon source. When reaching a cell density of 106 cells/mL, 0.8% galactose was added to induce the expression of Bax and trBcl-2like proteins. The expression was done overnight (16 hours). Mitochondria isolation, western-blot analyses on whole cell extracts and isolated mitochondria were done as described previously ^3333^.

### Statistical analysis

Error bars displayed on graphs represent the means ± S.E.M. of three independent experiments. Statistical significance was analyzed using the Student t-test. *P* < 0.05 was considered significant.

## ACKNOWLEDGEMENTS

We would like to thank Gabriel Ichim and Vincent Daubin for all the fruitful discussions. We would like to thank Laurent Coudert, Tim Hoffman, Kristina Radkova and Claire Céré for their technical assistance. This work was supported by Ligue nationale contre le cancer (Comité du Rhône).

## AUTHOR CONTRIBUTIONS

N.P performed all the experiments except for the following: L.J performed the cloning and the characterization of the trBcl-2L1 mutants. NP, T.T.M.N and N.R. performed the subcellular localization experiments. R.G., performed the cloning and expression of the codon optimized trBax sequence and deltaTM mutants as well as the cytosolic and mitochondrial calcium measurements. S.M., performed the yeast experiments; H.O. performed Trichoplax DNA/RNA extractions. N.P., H.O., B.S., R.R. and GG participate in the study design and writing the manuscript.

---

**Supplementary Figure S1.**
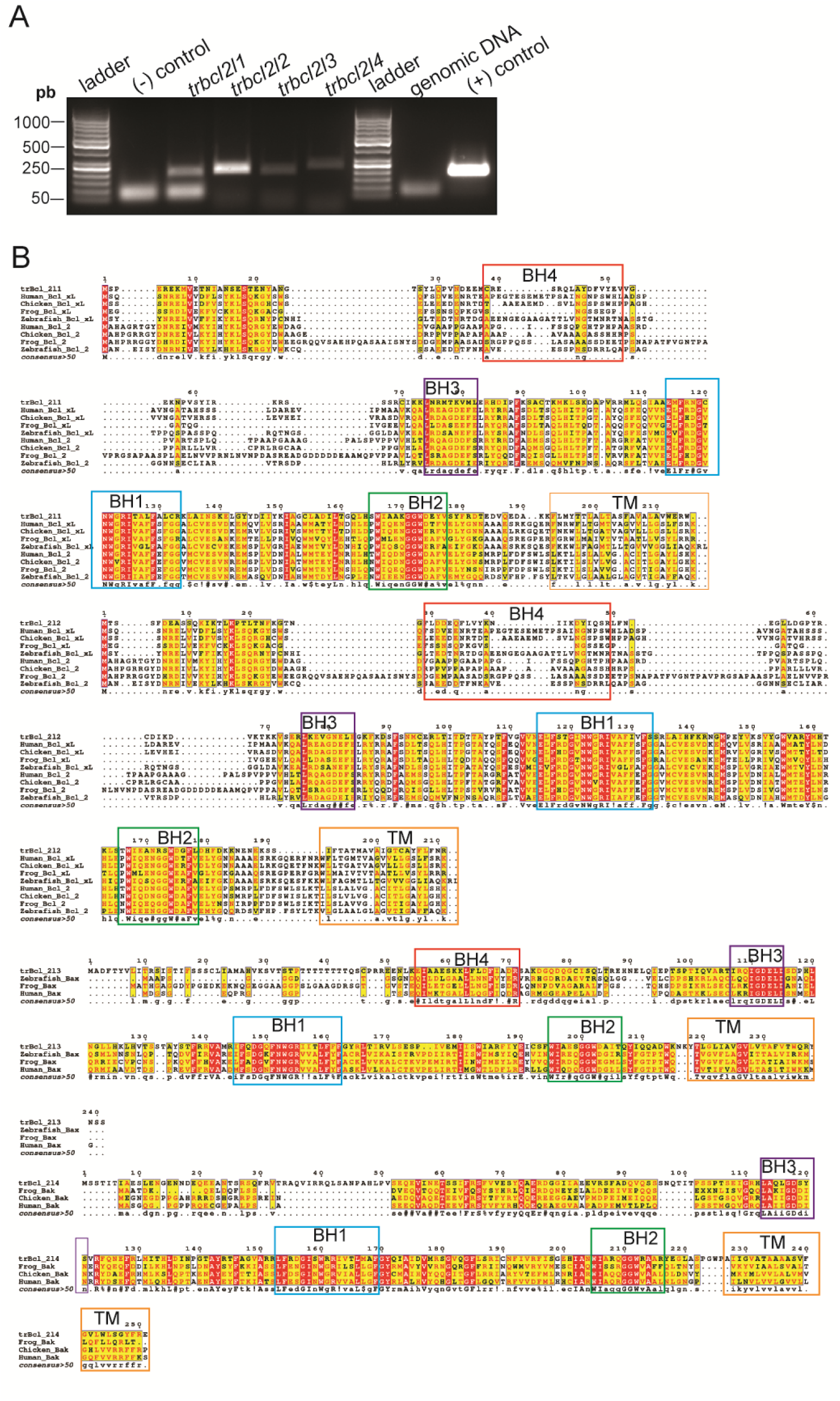
Expression and analysis of trBcl-2-like protein primary structures. **(A)** RT-PCR on RNA extracted from whole Trichoplax animals. *TrBcl2like* genes expression was detected using specific exon internal primers. DNA negative (- control) as well as genomic DNA were used as negative controls. L1 primers amplifying plasmid DNA encoding trbcl2l1 ORF was used as a positive PCR control. **(B)** ClustalW alignments of trBcl-2-like proteins with characterized Bcl-2 homologs in multiple species. Sequences corresponding to the BH homology and TM domains were highlighted with colored boxes.

**Supplementary Figure S2.**
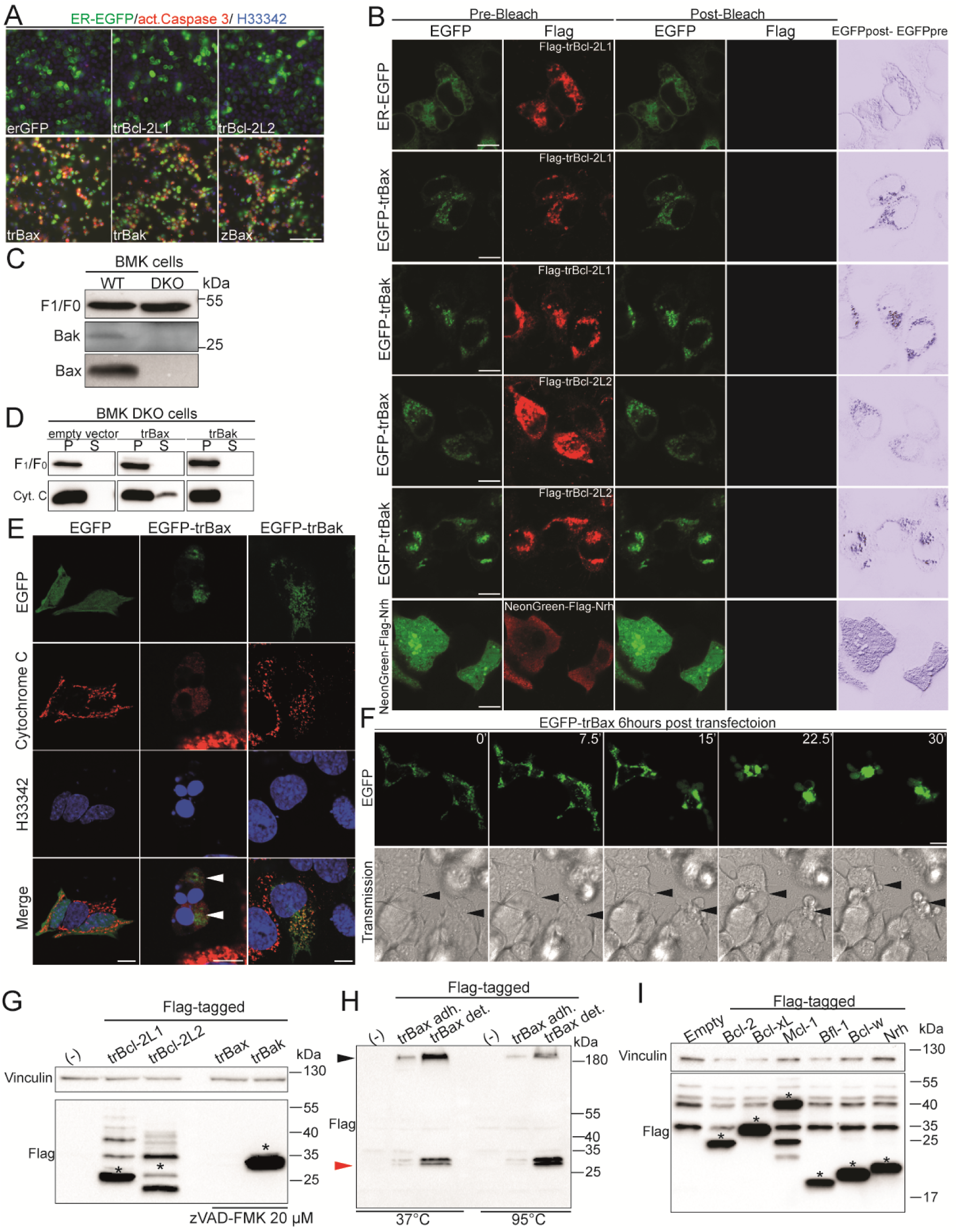
Control of MOMP by TrBcl-2-like proteins. **(A)** Representative florescence images of the effect of trBcl-2-like protein expression on Caspase 3 activation in HeLa cells. Cells were stained with anti-activated Caspase 3 antibody and counterstained with a nuclear dye H33342. mCherry and the zebrafish ortholog of Bax (zBax) were used as negative and positive controls, respectively. Scale bar: 100 µm. **(B)** Representative confocal images demonstrating the changes in EGFP (acceptor) fluorescence following Alexafluor 568 (donor) photobleaching. BMK cells expressing different combinations of EGFP-tagged trBax and Bak and Flag tagged trBcl-2L1 and trBcl-2L2 were fixed and immunostained with anti-Flag primary and anti-mouse IgG secondary antibodies linked with the Alexa568 fluorophore. Subtraction between EGFP fluorescence before and after photobleaching are shown in the right-hand column. NeonGreen-Flag-Nrh protein was used as positive control. Scale bar: 10 µm. **(C)** Immunoblot blot showing the content of endogenous Bax and Bak proteins in BMK WT versus BMK double *bax -/-, bak-/-* knock out cells. F_1_F_0_ ATPase was used as a loading control. **(D)** Immunoblot presenting the results of Cytochrome C release assay performed in BMK DKO cells. Cells transfected with empty vector or vector expressing trBax or trBak were subjected to Digitonin treatment (25 µg/mL) for 5 min. Cells were then separated by centrifugation in a pellet (P) fraction containing mitochondrial membranes, and supernatant (S) containing soluble proteins. The two fractions were immunostained with anti-Cytochrome C and anti-F_1_F_0_ ATPase antibodies. **(E)** Representative confocal images showing the subcellular distribution of Cytochrome C in EGFP-, EGFPtrBax- and EGFPtrBak-expressing BMK DKO cells. In control EGFPtrBak cells, Cytochrome C has a mitochondrial localization and becomes cytosolic following trBax induced MOMP. Scale bar: 20 µm. **(F)** Representative confocal images showing the changes in morphology of EGFPtrBax-expressing BMK cells. Following translation trBax was rapidly translocated to the mitochondria where it led to rapid cell shrinkage and cell blebbing. Scale bar: 10 µm. **(G)** Western blot (50 µg total proteins) showing the expression of trBcl-2-like proteins in BMK DKO cells. Asterisks highlight the bands corresponding to predicted molecular weights. Loading 50 µg of total protein extract and the use of zVAD-FMK (20 µM) was not sufficient to detect trBax. **(H)** Western blot (100 µg total proteins) of protein extracts obtained from detached cells and adherent cells expressing Flag-tagged trBax or transfected with empty vector (-). TrBax protein was highly enriched in the detached cells where it was detected as a multiprotein complex (black arrowhead) resistant to Laemmli treatment at 37°C but became monomeric (red arrowhead) once the temperature was raised to 95°C for 5 min. **(I)** Western blot analysis of the expression of Flag-tagged human Bcl-2 homologs BMK cells. The Vinculin protein was used as loading control.

**Supplementary Figure S3.**
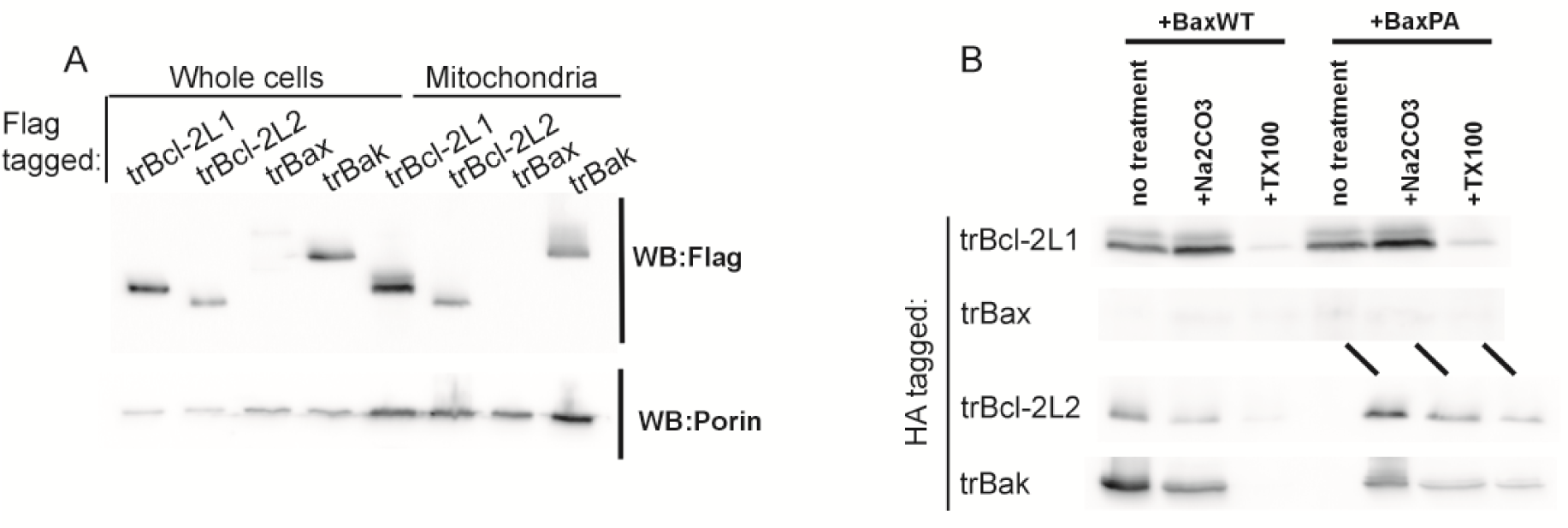
Analysis of trBcl-2like protein localization in the yeast model. **(A)** Western-blot analysis of trBcl-2like proteins expression in yeast mitochondria. Whole cell extracts (from ∼200,000 cells) or isolated mitochondria (50µg protein) were loaded on SDS-PAGE, blotted, and analyzed with anti-Flag (mouse monoclonal, Sigma) anti-yeast porin (mouse monoclonal, Thermofisher) antibodies. **(B)** Mitochondria isolated from strains coexpressing BaxWT or BaxP168A and the different trBcl-2like proteins were suspended (1mg/mL) in 0.6M mannitol, 2mM EGTA, 10mM tris/maleate, pH 7.0. 0.1mL of the same buffer, or of 1M Na2CO3 (pH 10.0) or of 1% Triton X100 were added and samples were incubated for 10 minutes on ice. They were centrifuged (10 minutes, 27,000xg), and pellets were analyzed by western-blotting for the presence of trBcl-2like proteins.

**Supplementary Figure S4.**
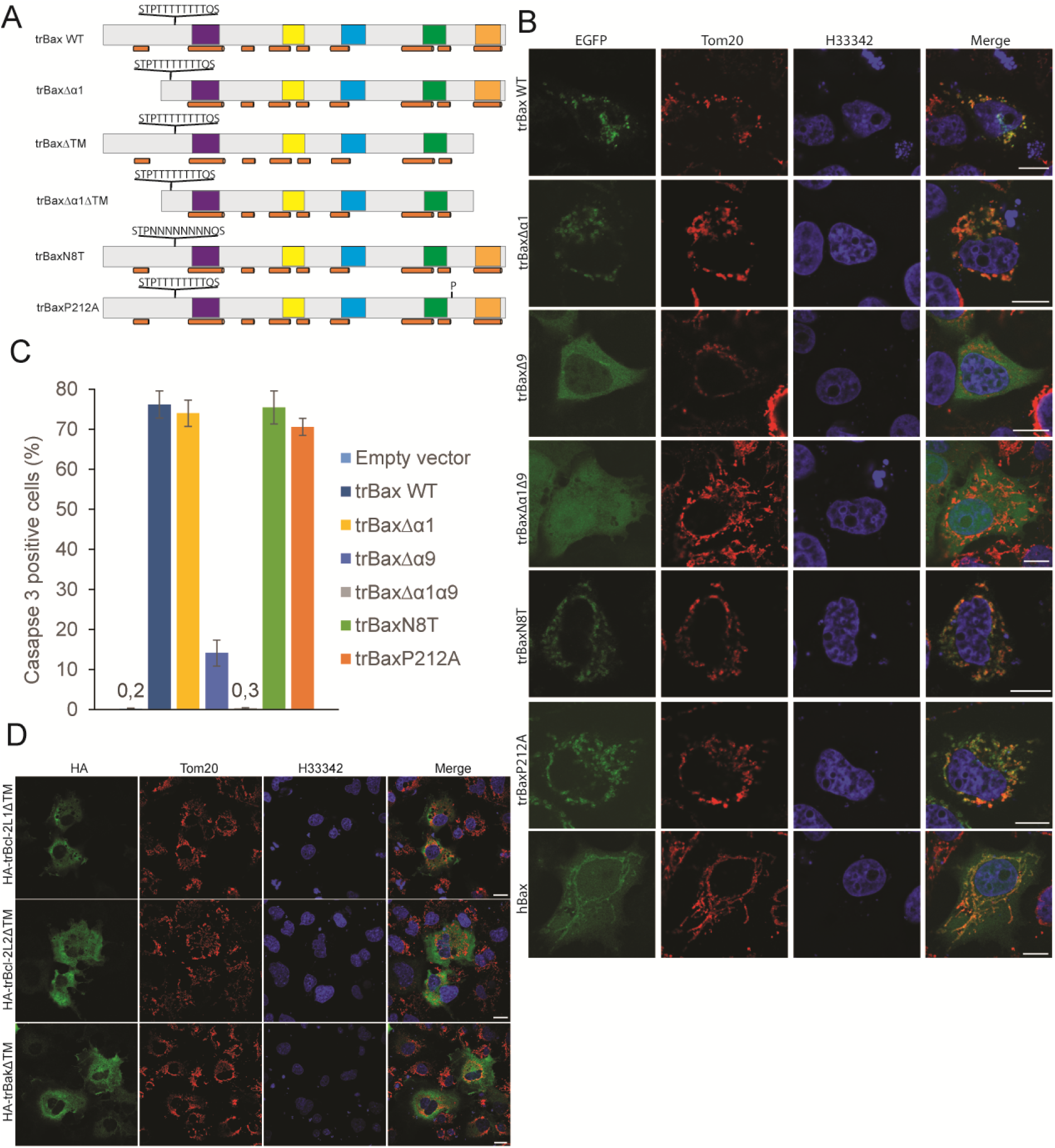
Subcellular localization of trBax dedicated mutants. **(A)** Schematic representation of the primary structures of trBax WT and associated mutants. The positions of conserved BH1-BH4 domains and of the C-terminus transmembrane domain (TM) as well as the locations of the predicted alpha helices are indicated with colored boxes and brown cylinders, respectively. **(B)** Representative confocal images showing the subcellular localization of EGFP-tagged trBax WT protein and associated mutants. Mitochondria were labeled using anti-Tom20 antibody. Merged channels between EGFP and Tom20 were presented on the right. Scale bar: 20 µm. **(C)** Histogram showing the effect of expression of trBax WT and associated mutants on Caspase 3 activation in BMK DKO cells. Deletion of the TM domain strongly impairs the pro-apoptotic activity of trBax (mean ± SEM; three independent experiments). **(D)** Representative confocal images showing the subcellular localization of HA-tagged trBcl-2-like ΔTM mutants. Mitochondria were labeled using anti-Tom20 antibody. HA fusion proteins were detected using anti-HA antibody. Merged channels between HA and Tom20 were presented on the right. Scale bar: 20 µm.

**Supplementary Figure S5.**
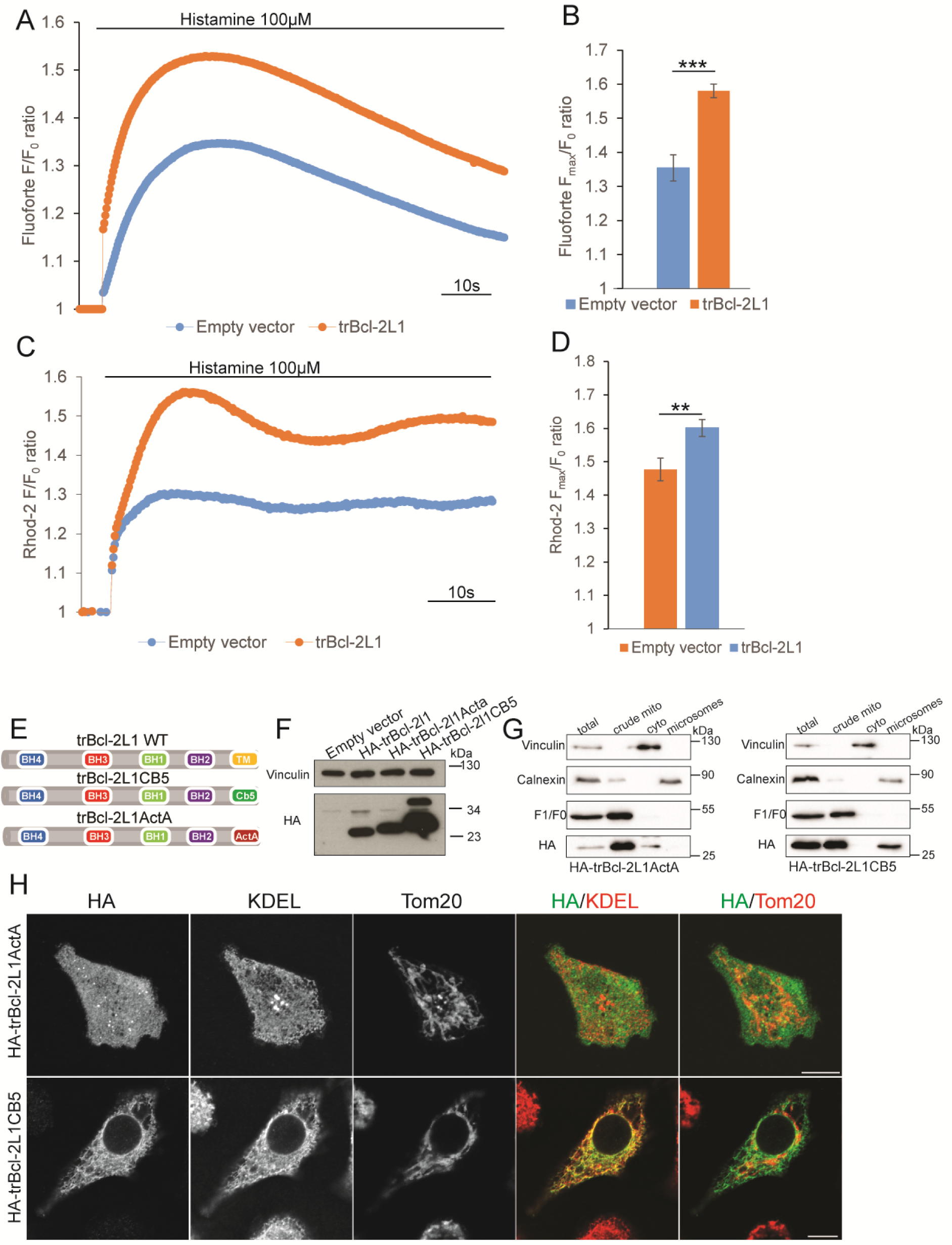
Effect of trBcl-2L1 expression on cytosolic and mitochondrial calcium fluxes. **(A)** Representative curve of cytosolic Ca^2+^ rise following ER-Ca2^+^ release induced by histamine (100 μM) stimulation. TrBcl-2L1 expression led to an increase in the ER-dependent cytosolic Ca^2+^ rise. Fluoforte fluorescence intensities were normalized against baseline values (F/F_0_ ratio). **(B)** Histogram depicting the maximum Ca^2+^ release in the cytosol (F_max_/F_0_ ratio) of HeLa cells following histamine (100 μM) treatment, from (E). (mean±S.E.M; n = 3 independent experiments). *** *P* < 0.001. **(C)** Representative curves of the relative changes in the mitochondrial Rhod-2AM fluorescence intensity normalized to the baseline (F/F_0_ ratio) reflecting the mitochondrial Ca^2+^ rise following histamine (100 µM) stimulation in HeLa cells expressing or not trBcl-2L1 protein. **(D)** Histogram depicting the maximum mitochondrial Ca^2+^ increase (F_max_/F_0_ ratio) following histamine (100 µM) treatment relative to control. (mean±S.E.M; three independent experiments). ** *P* < 0.01. **(E)** Schematic diagram presenting the construction of trBcl-2L1 mutants with single subcellular localization. BH and TM domains are presented with colored boxes. **(F)** Western blot showing the expression of HA-tagged trBcl-2L1 mutants with single subcellular localization in HeLa cells. **(G)** Subcellular fractionation of HeLa cells expressing HA-tagged trBcl-2L1ActA and CB5 mutants. Anti F_1_F_0_ ATPase, Calnexin and Vinculin antibodies were used as mitochondrial, ER and cytosol markers, respectively. **(H)** Representative confocal images of HeLa cells expressing trBcl-2L1 mutants with single subcellular localization. Subcellular localization of each protein was analyzed using anti-KDEL antibody for the staining of ER membranes anti-Tom20 antibody for labeling mitochondria anti-HA antibody for detection of HA tagged Bcl-2-like proteins. Merged channels between HA and KDEL and between HA and mitochondria are presented on the right. Scale bar: 10 µm

